# Defective Autophagy in *Sf1* Neurons Perturbs the Metabolic Response to Fasting and Causes Mitochondrial Dysfunction

**DOI:** 10.1101/2020.11.02.348789

**Authors:** Bérengère Coupé, Corinne Leloup, Julien Maillard, Luc Pénicaud, Tamas L. Horvath, Sebastien G. Bouret

## Abstract

**Objective:** The ventromedial nucleus of the hypothalamus (VMH) is a critical component of the forebrain pathways that regulate energy homeostasis. It also plays an important role in the metabolic response to fasting. However, the mechanisms contributing to these physiological processes remain elusive. Autophagy is an evolutionarily conserved mechanism that maintains cellular homeostasis by turning over cellular components and providing nutrients to the cells during starvation. Here we investigated the importance of the autophagy-related gene *Atg7* in *Sf1*-expressing neurons of the VMH in control and fasted conditions.

**Methods:** We generated *Sf1*-Cre; *Atg7*^loxP/loxP^ mice and examined their metabolic and cellular response to fasting.

**Results:** Fasting induces autophagy in the VMH, and mice lacking *Atg7* in *Sf1*-expressing neurons display altered regulation in glucose and leptin homeostasis and impaired energy expenditure regulation in response to fasting. Moreover, loss of *Atg7* in *Sf1* neurons causes alterations in the central response to fasting. Furthermore, alterations in mitochondria morphology and activity are observed in mutant mice.

**Conclusion:** Together, these data show that autophagy is nutritionally regulated in VMH neurons and that VMH autophagy participates in the control of energy homeostasis during fasting.

## 1. Introduction

Autophagy has received considerable attention over the past decade owing it to the fact that dysfunction of this cellular process, which degrades cytoplasmic materials including organelles and misfolded proteins, causes cellular alterations and contributes to a variety of diseases such as cancer, muscular disorders, and neurodegeneration [1–3]. Autophagy has also been implicated in physiological processes such as metabolic regulations. For example, the deletion of essential autophagy genes, such as the autophagy-related gene (Atg) 7, in the liver, pancreas, or adipose tissue produces alterations in body weight, adiposity, and glucose homeostasis [4–7]. More recent studies have shown the importance of autophagy in the central regulation of feeding behavior and energy homeostasis. Down-regulation of *Atg7* in the mediobasal hypothalamus increases energy intake and reduces energy expenditure resulting in weight gain and insulin resistance when animals are fed a high-fat diet (HFD) [8]. More specific conditional deletion of autophagy-related genes in pro-opiomelanocortin (POMC) and agouti-related peptide (AgRP) neurons revealed the importance of autophagy in the arcuate nucleus of the hypothalamus (ARH). Loss of *Atg7* in POMC neurons causes weight gain, hyperphagia, as well as leptin and insulin resistance [9–11]. Mice lacking *Atg7* in POMC neurons also exhibit increased sensitivity to weight gain when exposed to an HFD [10; 11]. Similarly, the loss of *Atg12* in POMC neurons promotes weight gain and perturbations in glucose homeostasis induced by HFD [12]. In contrast, deficiency in *Atg4* in POMC neurons or *Atg7* in AgRP neurons causes leanness, suggesting that autophagy induces distinct physiological effects based on the pathways and cell types engaged [13; 14].

Most of the research on hypothalamic autophagy has focused on neurons in the ARH thus far. However, although constitutive autophagy is also found in other hypothalamic nuclei, including in the ventromedial nucleus of the hypothalamus (VMH) [9], the importance of autophagy in VMH neurons has yet to be delineated. The VMH has long been associated with body weight regulation and glucose homeostasis. Physical lesions of the VMH induce hyperphagia, obesity, and diabetes [15]. Moreover, genetic studies have shown the importance of VMH neurons, particularly a subpopulation of VMH neurons expressing the steroidogenic factor-1 (SF-1), in relaying leptin and insulin action on energy balance and glucose metabolism [16–19]. In the present study, we examined the metabolic importance of autophagy in VMH neurons by generating mice that lack *Atg7* in *Sf1* neurons.

## 2. Material and methods

### 2.1 Animals

Mice were housed in individual cages under specific pathogen-free conditions, maintained in a temperature-controlled room with a 12 h light/dark cycle, and provided *ad libitum* access to water and standard laboratory chow (Special Diet Services). Animal studies were approved by the Institutional Ethics Committees of Care and Use of Experimental Animals of the University of Lille 2 (France). All experiments were performed in accordance with the guidelines for animal use specified by the European Union Council Directive of September 22, 2010 (2010/63/EU) and the approved protocol (APAFIS#13387–2017122712209790 v9) by the Ethical Committee of the French Ministry of Education and Research. Mice in which microtubule-associated protein one chain light chain (LC3) had been fused to the green fluorescent protein (GFP) were kindly provided by Dr. Mizushima [20]. To generate VMH-specific Atg7 knockout (*Sf1*-Cre; *Atg7*^loxP/loxP^) mice, *Sf1*-Cre mice (Jax mice stock# 006364) [16] were mated to mice carrying a loxP-flanked Atg7 allele (*Atg7*^loxP/loxP^) (C57BL/6 background, kindly provided by Dr. Komatsu) [21]. Breeding colonies were maintained by mating *Sf1*-Cre; *Atg7*^loxP/+^ mice to *Atg7*^loxP/loxP^ mice. All mice were generated in a C57BL/6 background. Animals were genotyped by PCR as described previously [16; 21]. Cre-negative *Atg7*^loxP/loxP^ were used as controls.

### 2.2. Fasting experiments

Mice were divided into the following groups: a first group of mice was fed standard chow *ad libitum* (also referred to as “fed” mice. A second group of mice was deprived of food for 12h (during the dark phase) before the experiment (also referred to as “fasted” mice). In some experiments, a third group of mice was deprived of food for 12h and then given free access to food for an additional 24h (also referred to as “refed” mice).

### 2.3. Physiological measures

One day after birth, the litter size was adjusted to 7 pups to ensure adequate and standardized nutrition until weaning. Male and female mice were weighed weekly from P7 through P119 using an analytical balance. Food intake, energy expenditure, and heat production were monitored in 2-3 months old mice using a combined indirect calorimetry system (TSE Systems). Briefly, O_2_ consumption and CO_2_ production were measured for 24 hours to determine the energy expenditure. In addition, food intake was determined continuously by the integration of weighing sensors fixed at the top of the cage from which the food containers were suspended into the sealed cage environment. Heat production was also measured through the experiment. At the end of the experiment, body composition analysis (fat/lean mass) was performed using an NMR (EchoMRI). These physiological measures were performed at the University of Cincinnati Mouse Metabolic Phenotyping Center. In addition, glucose and insulin tolerance were performed in 2-3 months old mice through the i.p. administration of glucose (1.5 mg/g body weight) or insulin (0.75 U/kg body weight) after overnight fasting, and the blood glucose levels will be measured 0, 15, 30, 45, 60, 90, 120, and 150 min following glucose challenge, as previously described [9]. Leptin sensitivity tests were performed in 2-3 months old male mice. Briefly, the mice were injected i.p. with vehicle (5 mM sodium citrate buffer) or leptin (3 mg/kg body weight, Peprotech) according to the following scheme: vehicle injections for five days, followed by leptin injections for three days. Body weight was measured during the injection period. Serum leptin and insulin levels were also assayed in 2-3 months old mice using leptin and insulin ELISA kits, respectively (Millipore).

### 2.4. Immunohistochemistry

Anesthetized male mice were perfused transcardially with 4% paraformaldehyde at 2-3 months of age. The brains were then frozen and sectioned at 30-um thick and processed for immunofluorescence using standard procedures. The primary antibodies used for IHC were as follows: rabbit anti-GFP (1:10,000, Invitrogen), rabbit anti-cFos (1:2,000, Oncogene), rabbit anti-p62/SQSTM1 (1:1,000, Abcam), rabbit anti-ubiquitin (1,000, Dako). The primary antibodies were visualized with Alexa Fluor 488 or Alexa Fluor 568 goat anti-rabbit IgGs (1:200, Invitrogen) and Alexa Fluor 568 donkey anti-sheep IgGs (1:200, Invitrogen). Sections were counterstained using bis-benzamide (1:10,000, Invitrogen) to visualize cell nuclei, and coverslipped with Fuoromount-G (SouthernBiotech).

### 2.5. Hypothalamic primary cultures

Hypothalami were collected from P2 wild-type mouse pups and dissociated by incubation with activated papain, followed by trituration accordingly to manufacturer protocol (Worthington). Dissociated cells were pelleted, then re-suspended in supplemented Neurobasal-A medium, and cultured onto cell culture slides coated with poly-L-lysine (Sigma-Aldrich). After one day *in vitro*, cells were transfected (T*rans*IT-TKO® Transfection Reagent, Mirus) with siRNA against Atg7 (50nM, Cell Signaling). Scramble siRNA (Cell Signaling) was used as control. After 72h, cells were incubated with 250nM MitoTracker® (Invitrogen), fixed in paraformaldehyde, and stained with MAP2 (1:500, Sigma). Experiments were independently repeated 3-4 times.

### 2.6. Image analysis

For the histological experiments, two sections through the VMH (for cFos, p62, and ubiquitin stainings and LC3-GFP fluorescence), the ARH and DMH (for cFos staining), and the ARH and PVH (for LC3-GFP fluorescence) from animals of each experimental group were acquired using a Zeiss LSM 710 confocal system equipped with a 20X objective. Slides were numerically coded to obscure the treatment group. Image analysis was performed using ImageJ analysis software (NIH) as described previously [9].

For the quantitative analysis of cell number, the number of cFos-immunopositive cells in the VMH, ARH, and DMH were manually counted. The average number of cells counted in two sections from each mouse was used for statistical comparisons.

For the quantification of ubiquitin- and p62-immunoreactivity and Mitotracker staining, each image plane was binarized to isolate labeled fibers from the background and compensate for differences in fluorescence intensity. The integrated intensity was then calculated for each image, which reflects the total number of pixels in the binarized image. This procedure was conducted on each image plane in the stack, and the values for all of the image planes in a stack were summed. The resulting value is an accurate index of the density of the processes in the volume sampled.

The NIH ImageJ macro called GFP-LC3 (http://imagejdocu.tudor.lu/author/rkd8/) [22] was used to quantify the number of LC3-GFP puncta.

### 2.7. Quantitative RT-PCR

VMH of 2-3 months old male mice fed *ad libitum* and fasted for 12h were microdissected. Total RNA was isolated using the Arcturus PicoPure RNA Isolation kit (Applied Biosystem). cDNAs were generated with the high capacity cDNA Reverse Transcription Kit (Applied Biosystem). Quantitative real-time PCR analysis was performed using the TaqMan Fast universal PCR Mastermix. mRNA expression was calculated using the 2-DDCt method after normalization with *Gapdh* as a housekeeping gene. Inventoried TaqMan® Gene expression assays *Atg5* (Mm00504340_m1), *Atg7* (Mm00512209_m1), *Atg12* (Mm00503201_m1), *InsR* (Mm01211875_m1), *Pomc (*Mm00435874_m1), *Npy (*Mm03048253_m1)*, Cpt1* (Mm01231183_m1), *Pgc-1α* (Mm01208835_m1), *Acaca (*Mm01304257_m1), *Ucp2* (Mm00627598_m1), and *Gapdh* (Mm99999915_g1), were used. All assays were performed using an Applied Biosystems Prism 7900HT fast sequence detection system.

### 2.8. Electron Microscopy

2-3 months old mice were perfused with 4% paraformaldehyde, and their brains were processed for electron microscopy examination. Briefly, ultrathin sections were then cut on a Leica ultra-microtome, collected on Formvar-coated single-slot grids, and analyzed with a Tecnai 12 Biotwin electron microscope (FEI).

### 2.9. Mitochondria activity

Mediobasal hypothalami from 2-3 months old male mice were dissected, and mitochondrial oxygen consumption was determined using high-resolution respirometry (Oxygraph-2K, Oroboros Instruments), as described previously [23]. Briefly, basal O_2_ flux of the respiratory chain was measured at 37 C without substrate or with a substrate for complex I (glutamate, 20 mM) and (D) complex II (succinate, 20 mM). States 3 and 4 respiration rates were achieved by adding to the incubation medium ADP (2.5 mM) and carboxy-atractyloside (CAtr, 0.75 μM), respectively. Measurement of maximal mitochondrial respiration was performed using Carbonyl cyanide m-chlorophenylhydrazone (CCCP, 0.5 μM), a chemical uncoupling molecule. Rotenone (0.5μM), a CI inhibitor, was used to measure the maximal respiration capacity due to complex II only. The respiratory control ratio (RCR) was calculated by dividing the state respiration 3 rate by the state 4 respiration rate. The experimental data were analyzed using DataGraph software (Oroboros Instruments).

### 2.10. Statistical analysis

All values were expressed as means ± SEM. Statistical analyses were conducted using GraphPad PRISM (version 5.0d). Statistical significance was determined using unpaired two-tailed Student’s t-tests, a one-way ANOVA followed by the Tukey post-hoc test, and a two-way ANOVA followed by the Bonferroni post-hoc test when appropriate. *P-*values less than 0.05 were considered as statistically significant.

## 3. Results

### 3.1. Fasting induces autophagy in the ventromedial nucleus of the hypothalamus

We previously reported that autophagy is constitutively activated in the hypothalamus of normal fed mice, including in the VMH [9]. However, how autophagy is regulated in specific hypothalamic nuclei remains largely unknown. To explore if nutrition influences autophagy in the VMH, we first examined the effect of food deprivation in transgenic mice in which microtubule-associated protein 1 light chain 3 (LC3) has been fused to the green fluorescent protein (GFP). Because LC3 is a reliable marker of autophagosomes, LC3-GFP mice can be used to monitor autophagy *in* vivo [24]. LC3-GFP puncta were readily detectable throughout the VMH of fed mice, and fasting markedly increased the density of LC3-GFP puncta found in both the dorsomedial and ventrolateral parts of the VMH (**Figure 1A** **and** **B**). Fasting-induced autophagy was not restricted to the VMH; the density of LC3-GFP puncta was also significantly higher in the ARH of fasted mice compared to fed mice and elevated in the PVH as well, although it did not reach statistical significance (**Figure S1A-B**). We next examined whether fasting caused changes in autophagy-related genes. Quantitative PCR analyses revealed that *Atg7* and *Atg12* mRNA levels were increased and decreased, respectively, in the VMH of fasted animals compared to fed mice (**Figure 1C**). However, mRNA expression of *Atg5* was unchanged (**Figure 1C**). Together, these data indicate that hypothalamic autophagy is nutritionally regulated and that fasting triggers autophagy in the VMH.

**Figure 1.**
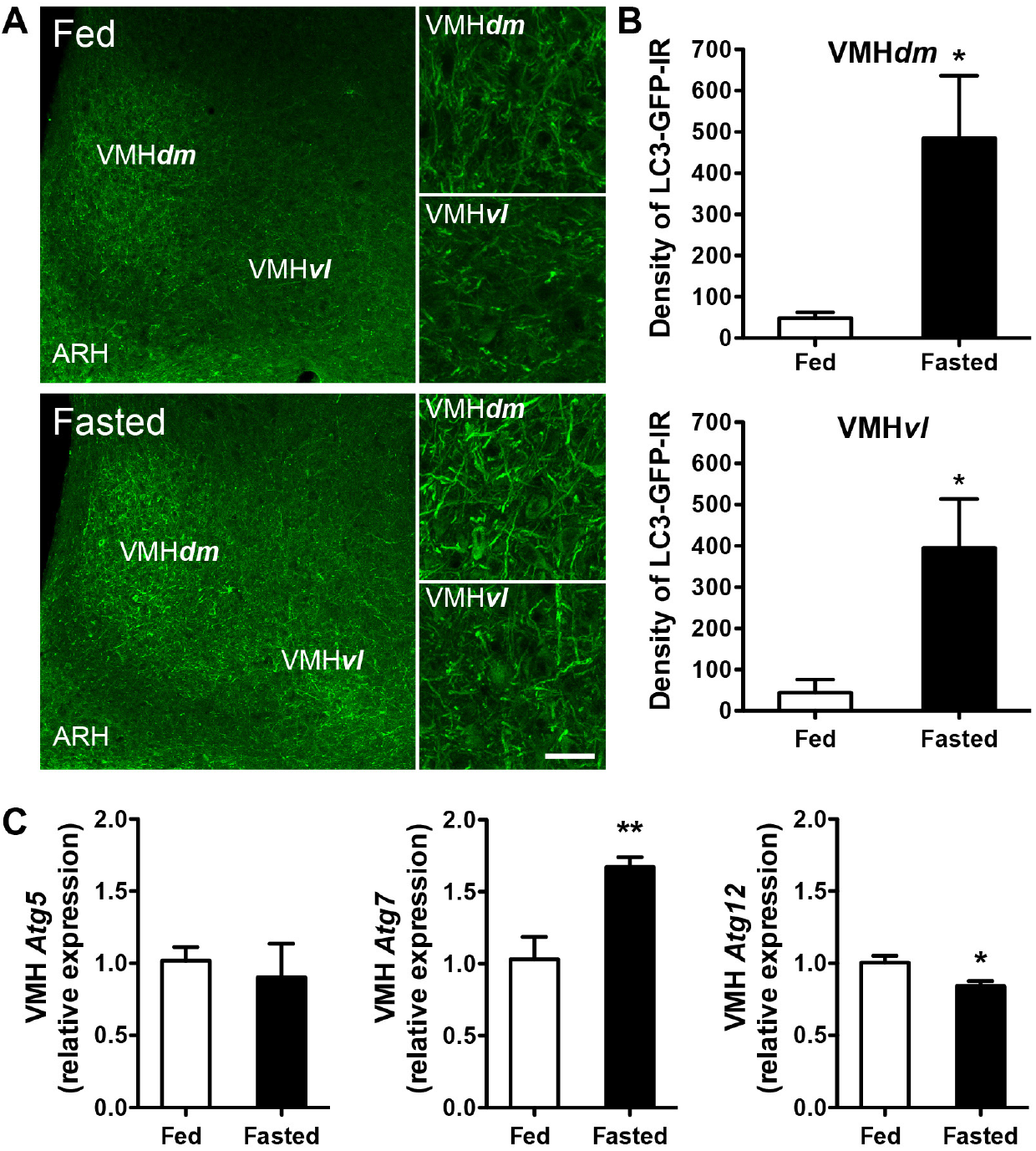
Fasting promotes autophagy in the ventromedial nucleus of the hypothalamus. (A) Representative images and (B) quantification of LC3-GFP immunofluorescence in the dorsomedial (dm) and ventrolateral (vl) parts of the ventromedial nucleus of the hypothalamus (VMH) of adult fed or 12h-fasted mice (n = 3 per group). (C) Autophagy-related gene mRNA levels in the VMH of adult fed or 12h-fasted mice (n = 3-5 per group). V3, third ventricle. Scale bar, 10 μm. Values are shown as mean ± SEM. \**P* ≤ 0.05, and ** *P* ≤ 0.01 *versus* fed.

### 3.2. Deletion of *Atg7* in SF1 neurons impairs metabolic response to fasting

To investigate the functional role of autophagy in VMH neurons, we generated mice lacking *Atg7* in cells that express *Sf1*, a factor predominantly expressed in VMH neurons (*Sf1*-Cre; *Atg7*^loxP/loxP^ mice) [16]. We first confirmed that the VMH is indeed autophagy-defective by analyzing the formation of ubiquitin- and polyubiquitin-binding protein p62/SQSTM1 because the accumulation of these aggregates is a hallmark of autophagy deficiency [25]. As expected, *Sf1*-Cre; *Atg7*^loxP/loxP^ mice displayed a 9- and 4-fold increase in the density of ubiquitin- and p62-immunoreactive aggregates, respectively, in the VMH (**Figure S2A** **and** **S2B**). Importantly, induction of ubiquitin and p62 immunoreactivity was not found in any other brain regions, confirming that the loss of autophagy in mutant mice occurs exclusively in the VMH and not in other brain sites expressing *Sf1*.

We next investigated the role of VMH autophagy in energy homeostasis and response to fasting. We first assessed food intake, locomotor activity, and energy expenditure (including O_2_ consumption, and CO_2_ and heat production) in fed and fasted mice. When fed *ad libitum*, *Sf1*-Cre; *Atg7*^loxP/loxP^ mice were comparable to their control littermate with regards to food intake (**Figure 2B**), locomotor activity (data not shown), oxygen consumption (VO_2_), carbon dioxide production (VCO_2_), and heat production (**Figures 2G, 2I, 2J, 2L** **and** **Figures S3A, S3C, S3D, S3F**). However, when fasted overnight, *Sf1*-Cre; *Atg7*^loxP/loxP^ mice displayed reduced VO_2_, VCO_2_, and heat production (**Figures 2H, 2I, 2K, 2L** **and** **Figures S3E, S3F**) as well as reduced cumulative food intake during a refeeding following an overnight fast (**Figure 2C**) compared to *Atg7*^loxP/loxP^ mice. However, this altered metabolic response was not accompanied by changes in carbohydrates *versus* lipid oxidation rates, as the RQ was normal in fed and fasted mutants (**Figures S3A-C**). Moreover, loss of *Atg7* in *Sf1* neurons did not result in changes in body weight (**Figure 2A**). Serum leptin levels were reduced in *Sf1*-Cre; *Atg7*^loxP/loxP^ mice (**Figure 2F**) despite an elevated fat mass (**Figure 2D**). Moreover, mutant mice showed an attenuated weight loss after exogenous leptin treatment (**Figure 2E**).

**Figure 2.**
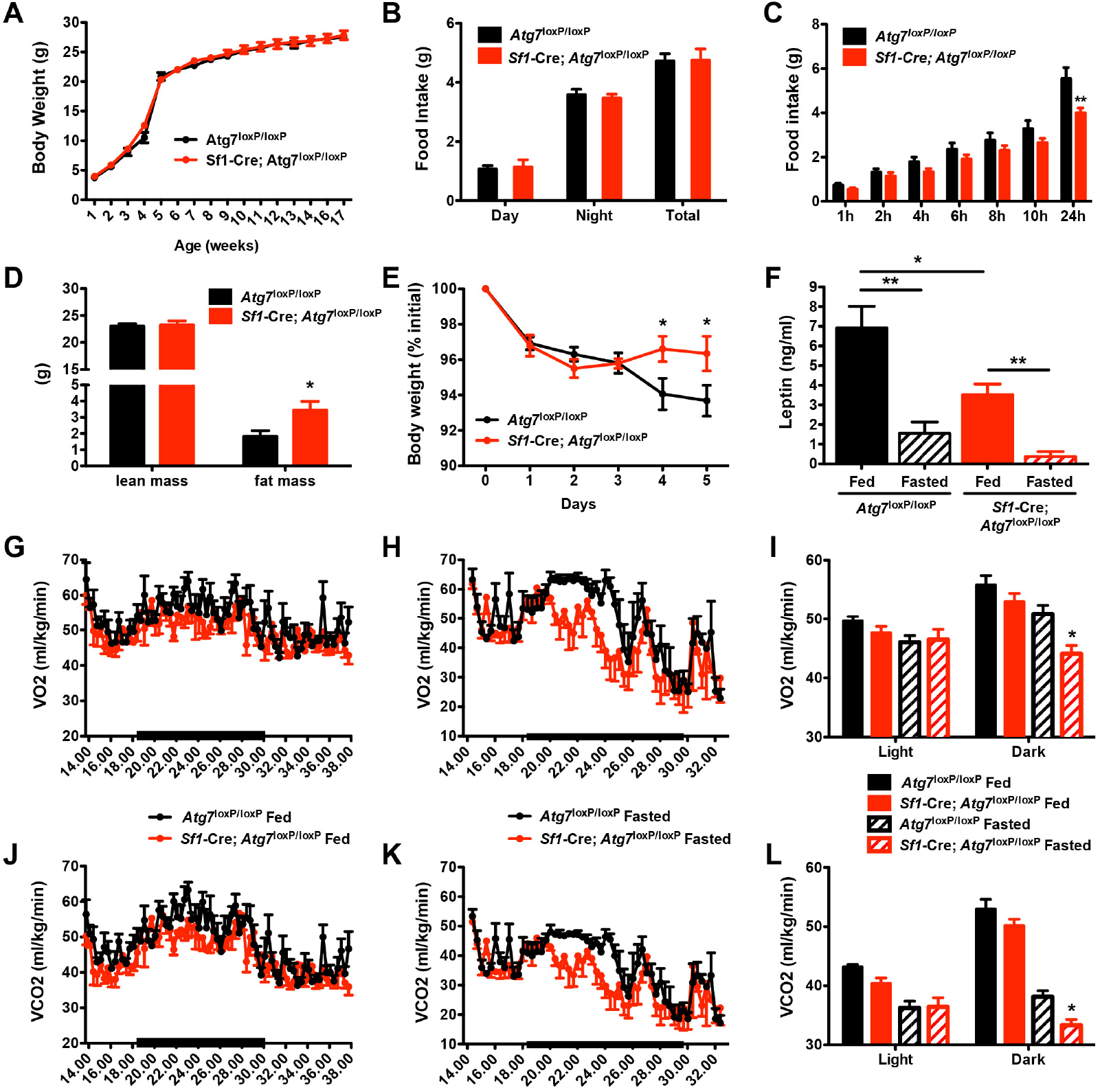
Altered energy homeostasis in mice lacking autophagy in SF1 neurons. (A) Body weight and (B) average food intake of *Atg7*^loxP/loxP^ and *Sf1*-Cre; *Atg7*^loxP/loxP^ (n 4-6 per group) male mice. (C) Food intake of refed of adult *Atg7*^loxP/loxP^ and *Sf1*-Cre; *Atg7*^loxP/loxP^ male mice after fasting (n = 5 for each group). (D) Body composition and (E) leptin sensitivity of adult *Atg7*^loxP/loxP^ and *Sf1*-Cre; *Atg7*^loxP/loxP^ male mice (n = 5-6 for each group). (F) Serum leptin levels in fed and 12h-fasted adult *Atg7*^loxP/loxP^ and *Sf1*-Cre; *Atg7*^loxP/loxP^ (n = 3-5 per group) male mice. (G-I) Oxygen (O2) consumption and (J-L) carbon dioxide (CO2) production in (G, J) fed and (H, K) 12h-fasted *Atg7*^loxP/loxP^ and *Sf1*-Cre; *Atg7*^loxP/loxP^ male mice (n = 5 for each group). Values are shown as mean ± SEM. \**P* ≤ 0.05, and *P* ≤ 0.01 *versus* each group.

Previous studies reported a role for SF1 neurons in glucose metabolism [16; 19; 26; 27]. We therefore measured several indices of glucose metabolism in *Sf1*-Cre; *Atg7*^loxP/loxP^ mice and found significantly lower fed plasma insulin concentrations compared to *Atg7*^loxP/loxP^ mice (**Figure 3B**) without changes in plasma glucose levels (**Figure 3A**). In addition, insulin receptor (*Insr*) mRNA levels in the VMH of fed *Sf1*-Cre; *Atg7*^loxP/loxP^ mice were significantly higher than those in control mice (**Figure 3C**). Fasting equally reduced glucose and insulin levels in *Sf1*-Cre; *Atg7*^loxP/loxP^ and *Atg7*^loxP/loxP^ mice (**Figure 3A-B**). We also performed glucose and insulin tolerance tests and found that *Sf1*-Cre; *Atg7*^loxP/loxP^ mice responded to a glucose challenge normally whereas they displayed insulin intolerance (**Figure 3D-E**).

**Figure 3.**
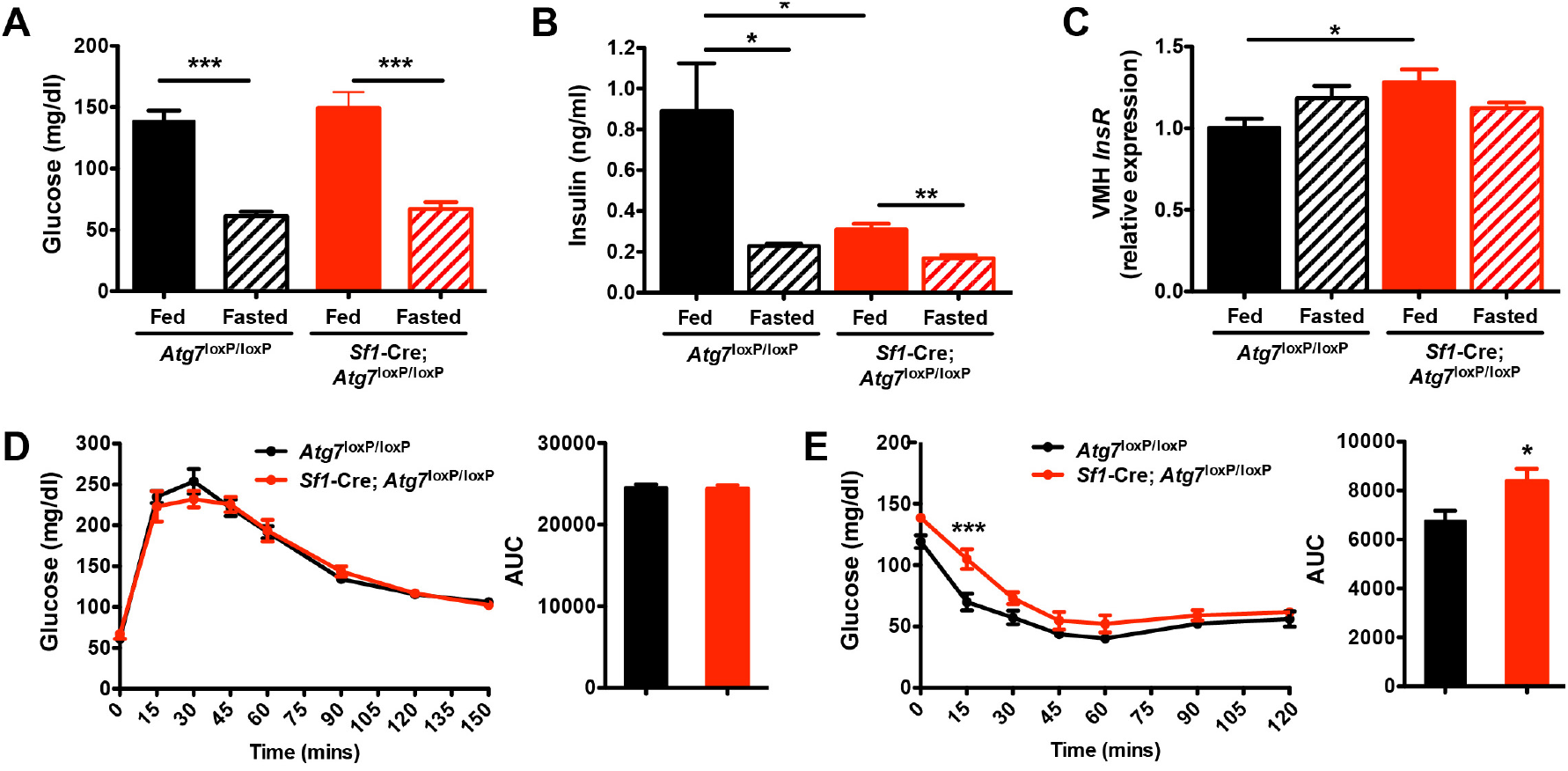
Glucose metabolism in *Sf1*-Cre; *Atg7*^loxP/loxP^ mice. (A) Serum glucose and (B) insulin levels, and (C) VMH insulin receptor (InsR) mRNA expression in fed and 12h-fasted adult *Atg7*^loxP/loxP^ and *Sf1*-Cre; *Atg7*^loxP/loxP^ (n = 4-7 per group) male mice. (D) Glucose and (E) insulin tolerance tests and area under the curves of adult *Atg7*^loxP/loxP^ and *Sf1*-Cre; *Atg7*^loxP/loxP^ (n = 4-6 per group) male mice. Values are shown as mean ± SEM. \**P* ≤ 0.05, \*\**P* ≤ 0.01, and \*\**P* ≤ 0.001 *versus* each group.

### 3.3. Altered hypothalamic response to fasting in *Sf1*-Cre; *Atg7*^loxP/loxP^ mice

The data described above suggest impairments in the physiological response to fasting in *Sf1*-Cre; *Atg7*^loxP/loxP^ mice. To determine whether these impairments are centrally mediated, we evaluated the ability of fasting to induce cFos expression (a surrogate marker of neuronal activation) in *Sf1*-Cre; *Atg7*^loxP/loxP^ and *Atg7*^loxP/loxP^ mice. Fed control and mutant mice had only a low density of cFos-immunoreactive cells in the ventromedial, arcuate (ARH), and dorsomedial (DMH) nuclei of the hypothalamus (**Figure 4A-C**). Fasting caused a 2-to-3-fold increase in the density of cFos-immunoreactive cells in the VMH, ARH, and DMH of *Atg7*^loxP/loxP^ mice; however, this induction is abrogated in *Sf1*-Cre; *Atg7*^loxP/loxP^ mice (**Figure 4A-C**). VMH neurons are predominantly glutamatergic and express synaptic vesicular transporter 2 (VGLUT2) [26]. *Sf1*-Cre; *Atg7*^loxP/loxP^ mice displayed lower *Vglut2* mRNA levels in the VMH than control mice, but *Vglut2* mRNA levels are unaffected by fasting in neither control and mutant mice (**Figure 4 D**). However, fasting induced down- and up-regulation of *Pomc* and *Npy* gene expression, respectively, in control mice, but this response is blunted in mutant mice (**Figure 4 E-F**).

**Figure 4.**
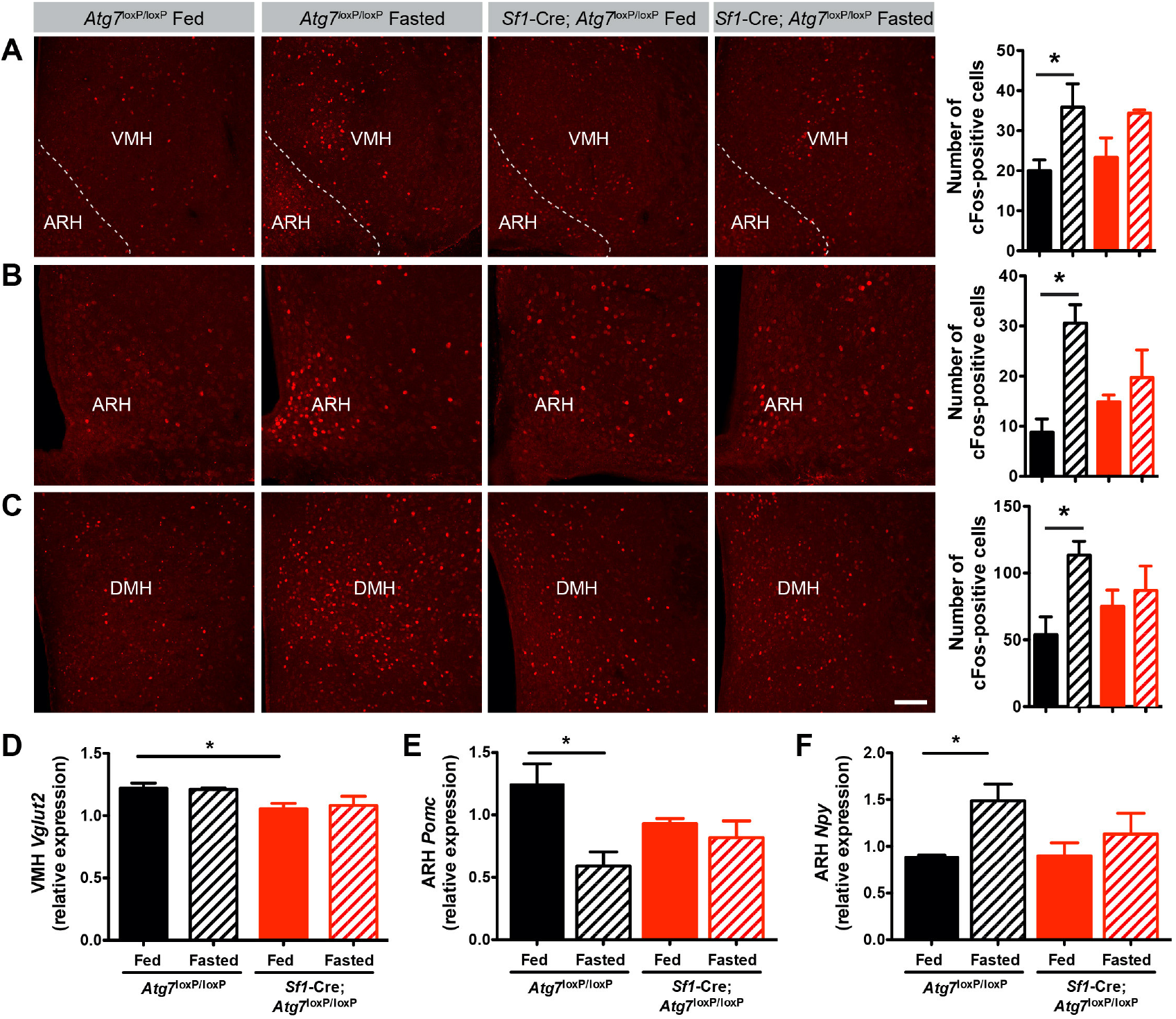
Fasting-induced hypothalamic neuronal activation is impaired in *Sf1*-Cre; *Atg7*^loxP/loxP^ mice. (A-C) Representative images and quantification of cFos-immunoreactive cells in the (A) ventromedial (VMH), (B) arcuate (ARH), and (C) dorsomedial (DMH) nuclei of the hypothalamus of fed and fasted adult *Atg7*^loxP/loxP^ and *Sf1*-Cre; *Atg7*^loxP/loxP^ male mice (n = 3-7 per group). (D) *Vglut2*, (E) *Pomc,* and (F) *Npy* mRNA expression in the ARH of adult fed and fasted *Atg7*^loxP/loxP^ and *Sf1*-Cre; *Atg7*^loxP/loxP^ male mice (n = 3-4 per group). Values are shown as mean ± SEM. \**P* ≤ 0.05 *versus* each group.

Because autophagy has been shown to maintain cell homeostasis and survival [21; 28], we also assessed whether *Atg7* deletion in *Sf1* neurons could affect the survival of these neurons using activated-caspase 3-immunoreactivity. The number of nuclei (stained by bisbenzimide) in the VMH of *Sf1*-Cre; *Atg7*^loxP/loxP^ mice was not different from that of *Atg7*^loxP/loxP^ mice (data not shown). We also counted the number of activated-caspase 3-positive cells (a surrogate marker of cell death), and we did not find any evidence of neuronal cell death (data not shown).

### 3.4. Loss of *Atg7* in VMH neurons alters mitochondrial morphology and function

Mitochondria play an important role in hypothalamic function [29–33] and autophagy is an important process for mitochondrial elimination [34–36]. We, therefore, investigated whether the deletion of *Atg7* in *Sf1* neurons causes mitochondria structural alterations using electron microscopy. Ultrastructural analysis showed a normal density of mitochondria related to cell size in VMH neurons of mutant mice (data not shown). However, there was a reduction in the number of mitochondria-autophagosome contacts in *Sf1*-Cre; *Atg7*^loxP/loxP^ mice compared to control mice (**Figure 5A**). Moreover, the relative aspect ratio, which is an indicator of mitochondrial length, was significantly increased in mutant mice compared to control mice (**Figure 5A**). To further examine the consequences of *Atg7* deficiency on mitochondria, we transfected primary cultures of hypothalamic cells with adenovirus containing siRNA directed against *Atg7* and stained mitochondria using MitoTracker. We found a 1.3-fold increase in the intensity of MitoTracker labeling *in Atg7* KD cells compared with control cells (**Figure 5B**). Collectively, these results indicate that both mitochondrial morphology and their association with autophagosomes are altered in the VMH of mutant mice.

**Figure 5.**
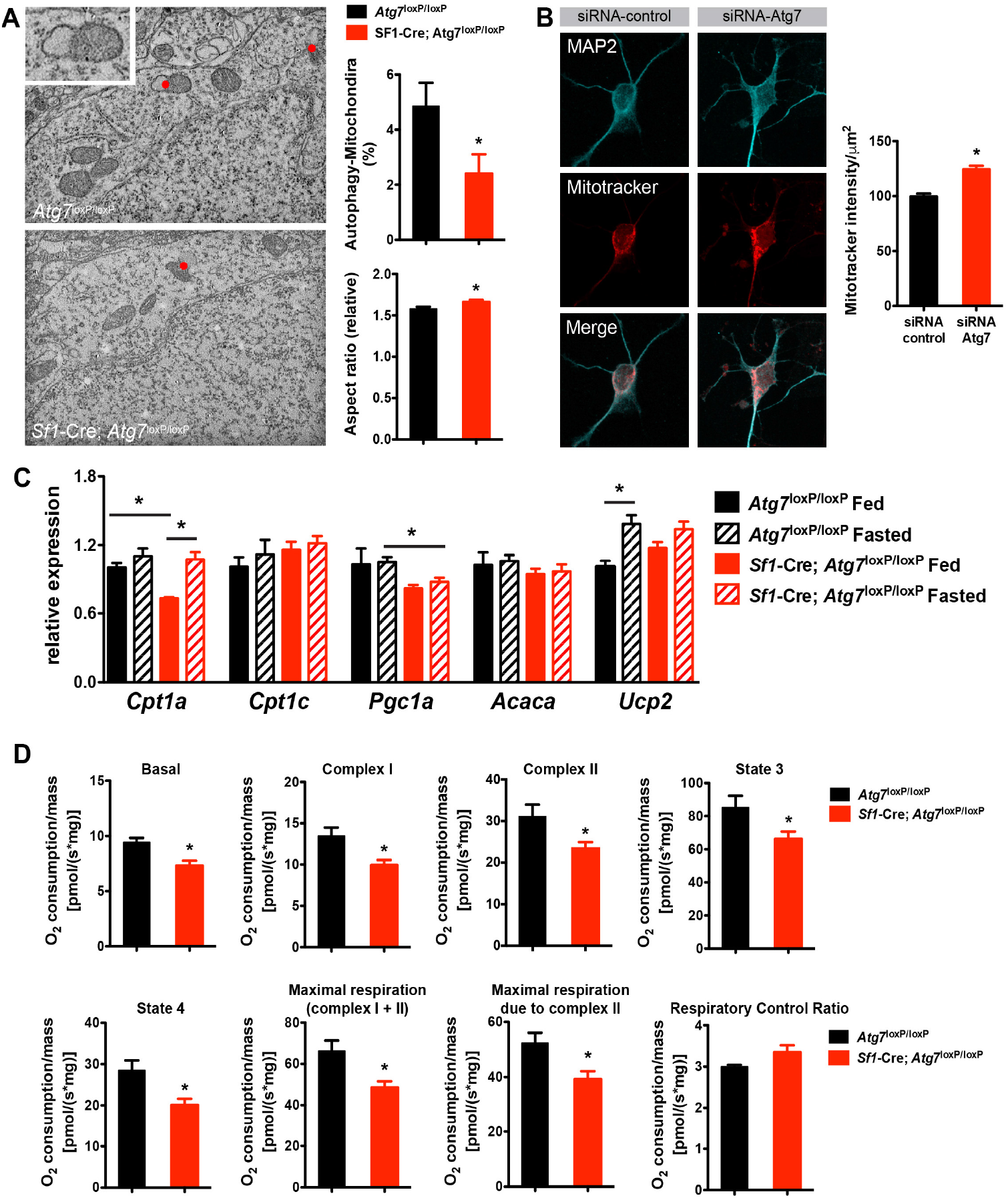
Loss of autophagy in SF1 neurons causes altered mitochondrial morphology and function. (A) Representative electron microscopy images of VMH neurons, percentage of autophagy-mitochondria contacts, and mitochondria aspect ratio in adult *Atg7*^loxP/loxP^ and *Sf1*-Cre; *Atg7*^loxP/loxP^ male mice (n= 4 per group). (B) Representative images and quantification of MitoTracker labeling in primary cultures of hypothalamic neurons 1 day after transfection with *Atg7* siRNA or scrambled siRNA (n = 3-4 per group). (C) Mitochondria-related gene mRNA levels in the VMH of adult (10-week-old) fed or 12h-fasted mice (n = 3-5 per group). (D) Measurement of mitochondrial respiration in the mediobasal hypothalamus of adult *Atg7*^loxP/loxP^ and *Sf1*-Cre; *Atg7*^loxP/loxP^ male mice (n = 4-8 per group). Values are shown as mean ± SEM. \**P* ≤ 0.05 *versus* each group.

We next measured mitochondria-related genes and found that *Cpt1a* and *Pgc1a* expression was downregulated in fed and fasted *Sf1*-Cre; *Atg7*^loxP/loxP^ mice, respectively, compared to controls (**Figure 5C**). Moreover, fasting induced *Ucp2* mRNA levels in the VMH of *Atg7*^loxP/loxP^ mice, but this response was blunted in mutant mice (**Figure 5C**). *Cpt1c* and *Acaca* mRNA levels were normal in both fed and fasted *Sf1*-Cre; *Atg7*^loxP/loxP^ and *Atg7*^loxP/loxP^ mice (**Figure 5C**). To explore the functional impact of *Atg7* deletion in *Sf1* neurons, we assessed mitochondrial respiration in the mediobasal hypothalamus of control and mutant mice. Basal oxygen consumption was approximately 20% lower in *Sf1*-Cre; *Atg7*^loxP/loxP^ mice compared to control animals (**Figure 5D**). A significant reduction in mitochondrial oxygen consumption was also observed after the addition of complex I (glutamate) and II (succinate) substrates (**Figure 5D**). Moreover, *Sf1*-Cre; *Atg7*^loxP/loxP^ mice displayed a reduction in coupled (state 3) and uncoupled (state 4) mitochondrial respiration (**Figure 5D**). CCCP-induced maximal respiration was also attenuated in mutant mice, and this effect appears unaffected by the complex I inhibitor rotenone (**Figure 5D**).

Together, these data indicate that mitochondria morphology and activity are altered in mice lacking *Atg7* in *Sf1* neurons.

## 4. Discussion

Autophagy has been involved in a variety of physiological and pathological pathways, and more recent studies demonstrated the importance of hypothalamic autophagy in energy balance regulation and glucose homeostasis. Here we show that in fed conditions, loss of the autophagy gene *Atg7* in *Sf1* VMH neurons has no effects on body weight, food intake, and energy expenditure, and only has a moderate impact on glucose metabolism. However, when fasted, *Sf1*-Cre; *Atg7*^loxP/loxP^ mice display abnormalities in O2consumption, CO2 and heat production, and they consume less food after refeeding. These findings differ from other studies showing that loss of Atg7 in arcuate POMC neurons increases body weight and fat mass, decreases O2 consumption, and impairs glucose metabolism [9–11]. These data indicate that the autophagy Atg7 pathway exerts distinct physiological functions depending on the neuronal cell type that is engaged. Consistent with this idea, mice lacking *Atg7* in AgRP neurons are leaner and display an opposite phenotype than mice lacking Atg7 in the neighboring POMC neurons [14].

The mediobasal hypothalamus, including the VMH, plays a critical role in sensing and integrating metabolic and nutrient signals from the periphery and engaging neural circuits to control feeding and glucose regulation. Autophagy is an essential component of the cellular response to starvation for energy provision [37]. In the present study we show that fasting triggers autophagy in the VMH as well as in other hypothalamic nuclei such as the ARH. These findings are in good agreement with a previous study from Kaushik and colleagues that showed that starvation increases LC3-II/I in the mediobasal hypothalamus, a region comprising both the arcuate and the ventromedial nuclei [14]. Leptin and insulin are likely candidates to mediate the effect of fasting on VMH autophagy. Fasting reduces leptin and insulin levels, and we recently reported that leptin-deficiency induces autophagy in the hypothalamus [38]. In addition, the VMH expresses high levels of leptin and insulin receptors [39; 40], and their loss in VMH Sf1 neurons influences the response to HFD-induced metabolic disturbances [16; 41]. Moreover, electrophysiological studies indicated the effects of leptin and insulin on VMH SF1 neurons is mediated through PI3K [42], which is a critical signaling pathway involved in autophagy activation [43]. It is likely that the lack of autophagy in *Sf1*-expressing neurons also indirectly impacts neurons outside of the VMH. For example, a previous study showed that POMC neurons in the arcuate nucleus receive strong excitatory inputs from the VMH and that the strength of the excitatory input from the VMH to POMC neurons is diminished by fasting [44]. Interestingly, in the present study, we show that mutant mice exhibit a reduction in *Vgltu2* expression associated with an impaired response of POMC neurons to fasting. However, different nutritional signals might trigger a distinct autophagy response. For example, HFD reduces autophagy in the hypothalamus, as evidenced by a reduction in Atg5, Atg7, and LC3-II/I expression [8]. Moreover, loss of Atg7 in the mediobasal hypothalamus or POMC neurons exacerbates the effects of HFD on adiposity and glucose homeostasis [8]. Although beyond the scope of this study, it would be interesting to examine whether HFD exerts similar or opposite effects in mice lacking Atg7 in VMH Sf1 neurons.

Neuronal function is energetically costly, and mitochondria almost exclusively provide this energy need. In addition, to produce energy in the form of adenosine triphosphate via oxidative phosphorylation, mitochondria also play many other important roles, such as buffering cytosolic Ca^2+^, synthesizing lipids, and producing signaling molecules such as reactive oxygen species (ROS) [45]. To carry out these functions effectively, mitochondria undergo morphological changes and localize to specific cellular compartments as required, brought about by fission and fusion. A series of studies have highlighted the importance of the hypothalamic mitochondria dynamic. Disruption of mitochondria fusion in arcuate AgRP neurons protects against HFD-induced obesity [32]. In contrast, it causes obesity when it is blocked in POMC neurons [31]. Moreover, dynamin-related protein1-dependent mitochondrial fission is a key mechanism in nutrient sensing in the ventromedial part of the hypothalamus [23]. It is also essential to have cellular processes in place to ensure the degradation of mitochondria that are damaged beyond repair, and this process is called mitophagy. In the present study, we report that loss of *Atg7* in *Sf1* neurons causes a reduction in mitochondria-autophagosome contacts, suggesting impairments in mitophagy in mutant mice. Notably, this abnormal mitophagy appears to be associated with alterations in mitochondria morphology and respiration. UCP2 has been linked with mitophagy [46], and VMH UCP2 expression regulates mitochondria fission, which in turn controls glucose regulation [47]. Consistent with these findings, we report here that *Ucp2* expression is dysregulated in *Sf1*-Cre; *Atg7*^loxP/loxP^ mice. Together, these observations suggest that the UCP2-autophagy pathway is a critical component of VMH mitochondria dynamic and function. In addition to play a role in mitochondrial dynamic, autophagy is also involved in the lipolytic process to break down cellular lipid stores and provide energy to cells, including hypothalamic neurons, in fasting conditions [14; 48; 49]. However, whether VMH autophagy controls mitochondria oxidative metabolism and avoids fatty acids toxicity in starved cells remains to be investigated.

## Author contributions

Conceptualization, B.C. and SGB; Methodology, BC, CL, JM, and TLH; Formal Analysis, BC, JM, CL, and TLH; Writing - Original Draft, BC; Writing - Review & Editing, SGB; Supervision, LP and SGB; All authors contributed to editing the manuscript and approved the manuscript for publication.

## Acknowledgments

We thank Drs. Masaaki Komatsu and Noboru Mizushima for the generous gift of the *Atg7*^loxP/loxP^ and LC3-GFP mice, respectively. We also would like to thank the University of Cincinnati Mouse Metabolic Phenotyping Center (supported by grant U24DK059630).

## Appendix A. Supplementary Data

Supplementary data to this article can be found online.

---

**Supplemental Figure 1.**
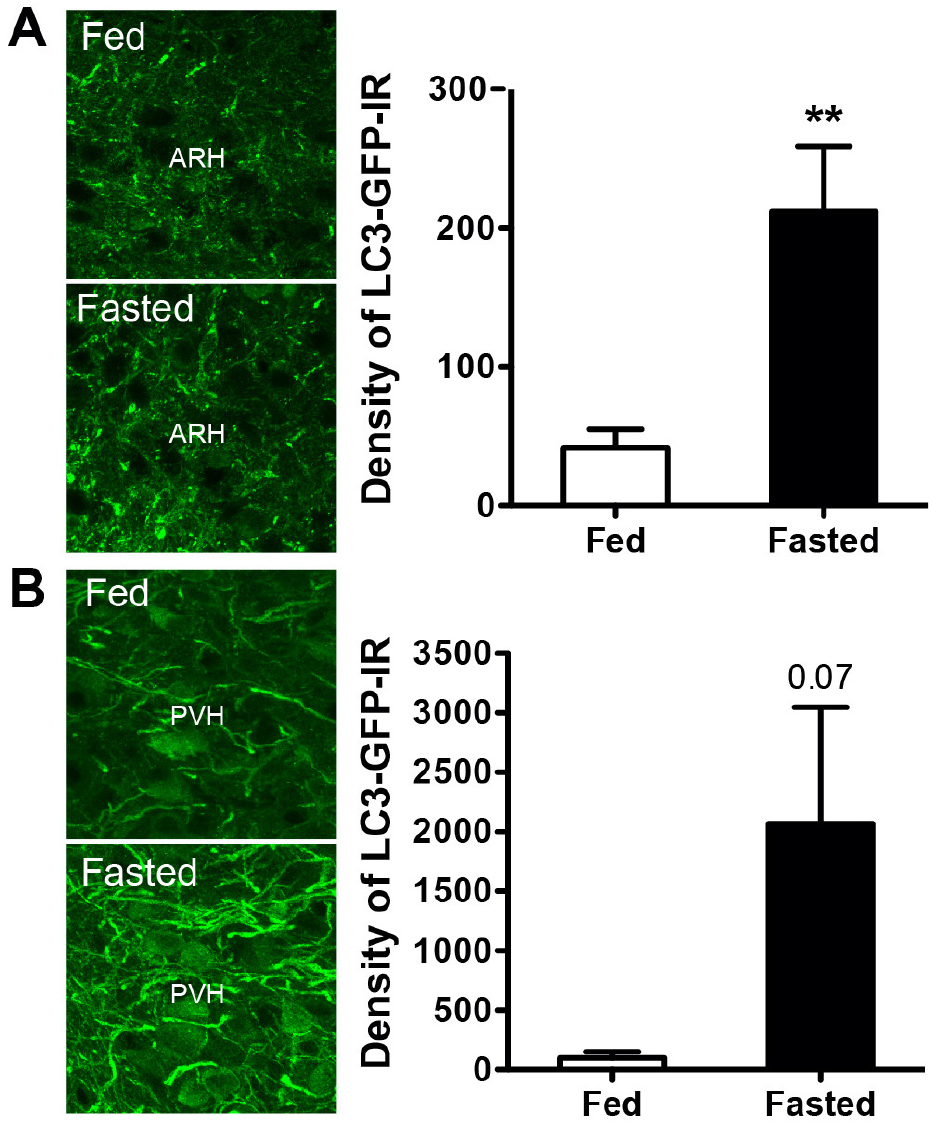
Fasting promotes autophagy in the arcuate nucleus of the hypothalamus. Representative images and quantification of LC3-GFP immunofluorescence in (A) the arcuate (ARH) and (B) paraventricular (PVH) nuclei of the hypothalamus of adult fed or 12h-fasted mice (n = 3-5 per group). Values are shown as mean ± SEM. \**P* ≤ 0.01 versus fed.

**Supplemental Figure 2.**
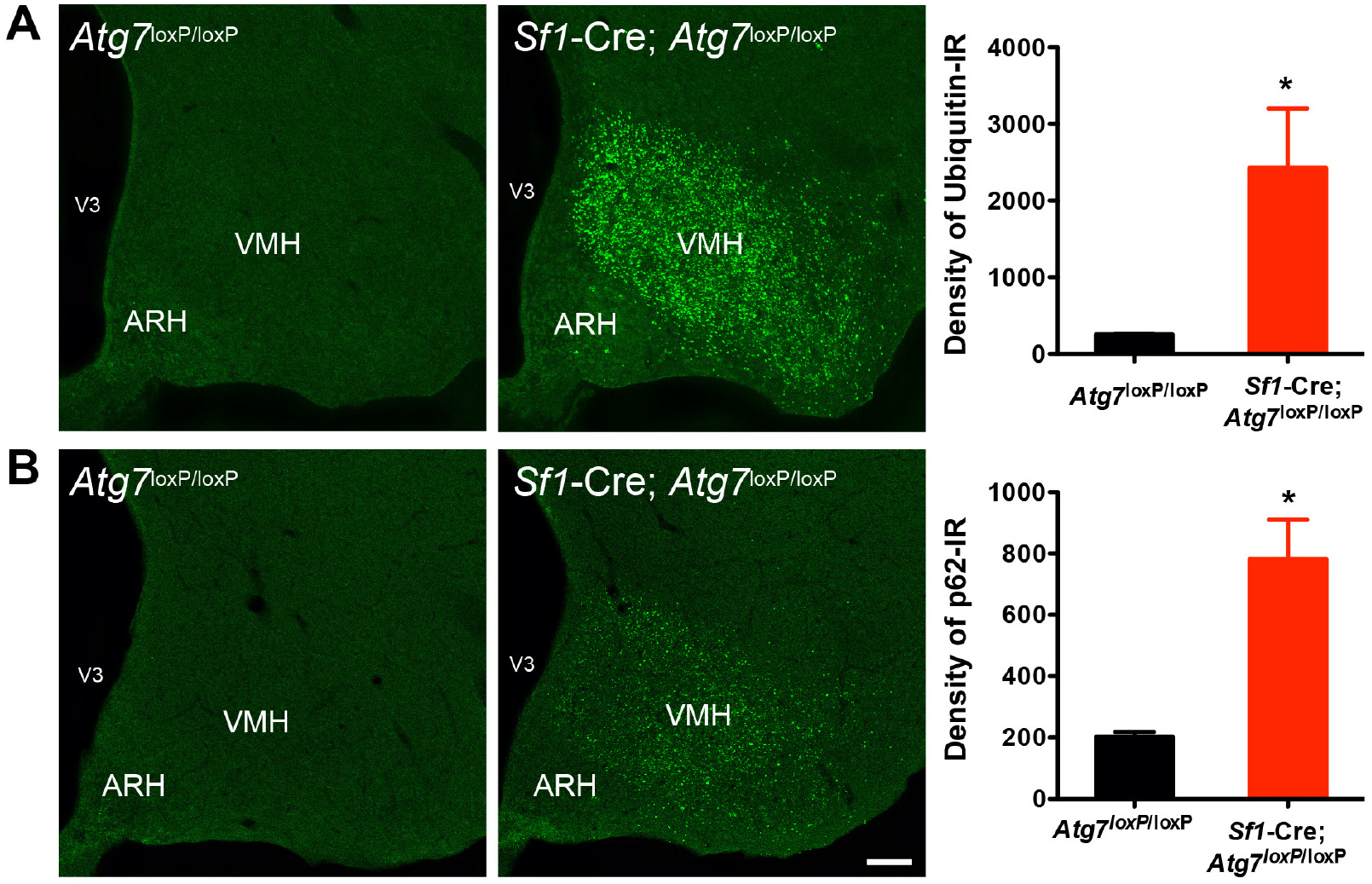
Loss of autophagy in SF1 neurons leads to the accumulation of ubiquitin aggregates in the ventromedial nucleus of the hypothalamus. Representative images and quantification of (A) ubiquitin- and (B) p62-immunoreactivity in the ventromedial nucleus of the hypothalamus (VMH) of adult *Atg7*^loxP/loxP^ and *Sf1*-Cre; *Atg7*^loxP/loxP^ male mice (n = 2-3 per group). ARH, arcuate nucleus of the hypothalamus; V3, third ventricle. Scale bar, 50 μm. Values are shown as mean ± SEM. \**P* ≤ 0.05 versus *Atg7*^loxP/loxP^.

**Supplemental Figure 3.**
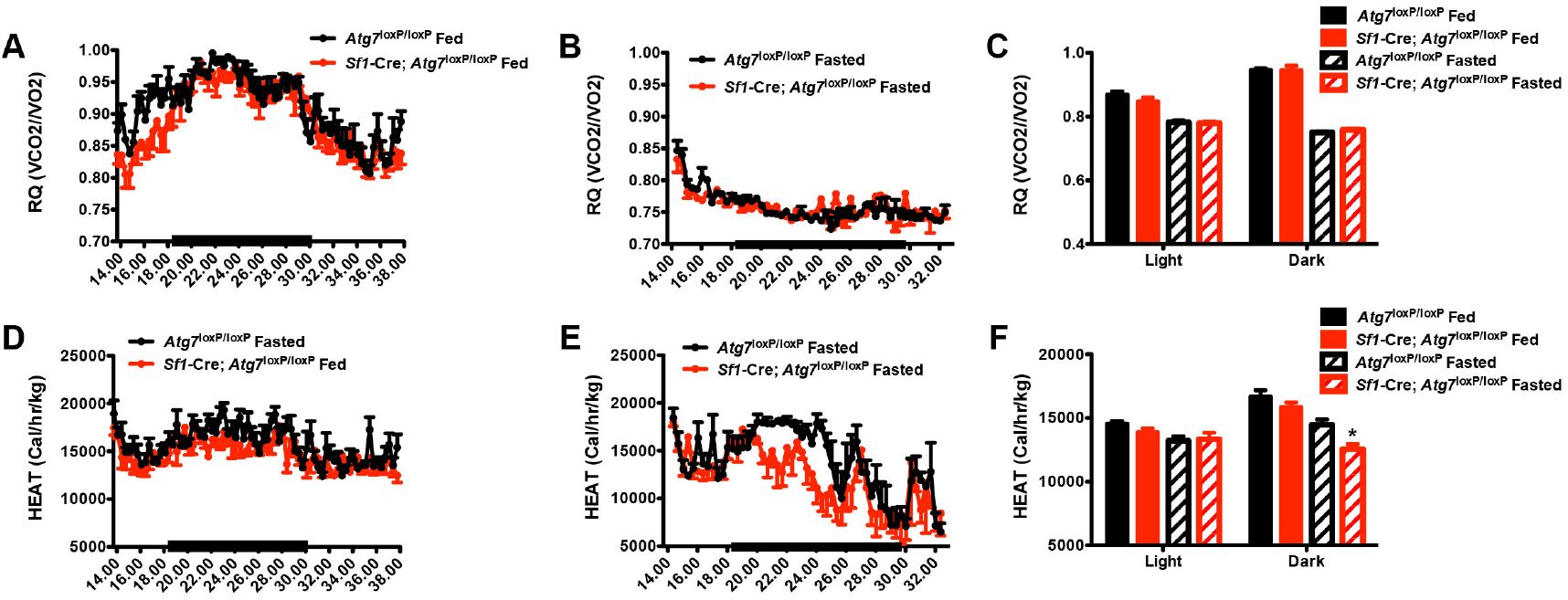
Altered heat production in mice lacking autophagy in *Sf1* neurons. (A-C) Respiratory quotient and (D-F) heat production in (A, D) fed and (B, E) 12h-fasted *Atg7*^loxP/loxP^ and *Sf1*-Cre; *Atg7*^loxP/loxP^ male mice (n = 4 for each group). Values are shown as mean ± SEM. \**P* ≤ 0.05 versus each group.

